# An integrated meta-omics approach for identifying candidate organic micropollutant degraders in complex microbial communities

**DOI:** 10.1101/2025.02.21.639455

**Authors:** Tal Elad, Kai Tang, Michaël Pierrelée, Marlene Mark Jensen, Mikkel Bentzon-Tilia, Barth F. Smets, Arnaud Dechesne, Borja Valverde-Pérez

## Abstract

Biotransformation is a significant determinant of the fate of organic micropollutants (OMPs) in natural and engineered environments; yet identifying OMP-transforming microorganisms remains challenging. Integrating metagenomics and metatranscriptomics, we searched for correlations between the biotransformation of ammonia and atenolol and the transcriptional activity of metagenome-assembled genomes (MAGs) across five nitrifier-rich cultures in batch experiments. The biotransformation of ammonia correlated with the activity of ammonia-oxidizing bacterium *Nitrosomonas europaea* but not with the activity of other bacteria, including several ammonia oxidizers. Additionally, the biotransformation of ammonia correlated with the transcript abundance of the ammonia monooxygenase (AMO) expressed by *N. europaea* but not with the transcript abundance of AMO at the community level. Atenolol biotransformation correlated with the activity of four MAGs representing three heterotrophic genera: *Terrimonas*, *Flavobacterium*, and *Zeimonas*. It did not correlate with the total transcript abundance of any member of a comprehensive set of amidohydrolases, which are predicted to transform this drug. By contrast, it correlated with the expression of the amidohydrolase asparagine synthase (AsnB) identified in the *Terrimonas* and *Flavobacterium* MAGs. In summary, we present a novel association-based method for elucidating biotransformation processes robust to variability in enzyme reaction kinetics with implications for OMP control.

**Synopsis:** The fate of organic micropollutants in natural and engineered systems depends greatly on biotransformation by microbes. This study proposes an approach for identifying these microbes with implications for the study and control of organic micropollutant removal.

**ABSTRACT GRAPHICS:** 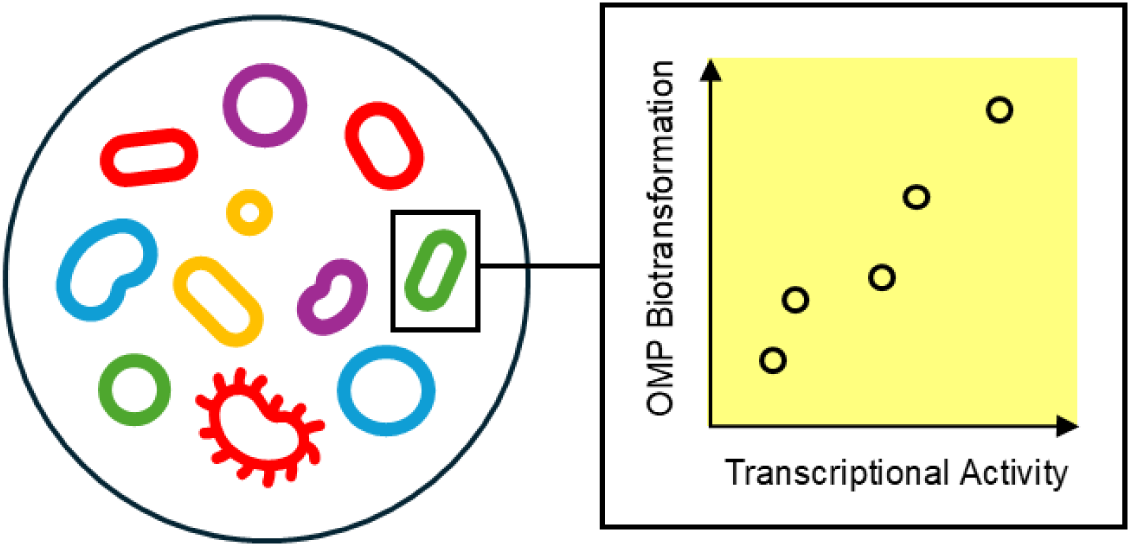

## INTRODUCTION

Pharmaceuticals, corrosion inhibitors, pesticides, and other organic chemicals used by modern society occur at trace concentrations (nanograms to micrograms per liter) in diverse aquatic systems.^1^ Potentially hazardous to human health and the environment, such chemicals raise increasing concerns and are generally regarded as organic micropollutants (OMPs).

Transformation by microorganisms is a significant determinant of OMP fate in natural and engineered systems^2–4^ and depends on environmental or process parameters.^5–7^ However, identifying OMP degraders remains challenging. A better understanding of which microbes degrade which OMPs and via which enzymatic pathways will facilitate the prediction and enhancement of OMP removal.

Currently, OMP degraders can be identified by stable isotope probing (SIP).^8–10^ SIP involves labelling a growth substrate with a stable isotope and tracking the isotope in biomolecules and cellular components of specific microbes. However, this approach has limitations. Firstly, OMPs can be transformed by enzymes with different primary substrates and low specificities, without being assimilated by the cells and contributing to growth.^11,12^ Secondly, the biotransformation of OMPs can proceed as a sequence of reactions that are not carried out by a single microbe. If so, the isotopes will not necessarily be assimilated by the cells involved in the first reaction, which determines the rate of removal. An alternative experimental method is enzyme inhibition; yet this method typically allows only for a crude distinction, e.g., between the contributions of ammonia-oxidizing and heterotrophic bacteria.^13–15^

Johnson *et al.* suggested to mine shotgun metatranscriptomic data for positive relationships between OMP biotransformation rate constants and the expression of specific genes by complex microbial communities to identify potential OMP-transforming enzymes.^16^ More recently, Achermann *et al.* identified positive relationships between oxygenases and oxidative biotransformations of OMPs by activated sludge within metatranscriptomics data.^17^ These metatranscriptomic association studies represent what can be considered an enzyme-centric approach, as they pool together all the enzymes with the same function in different microbes and implicitly assume they contribute equally to their substrate transformation. This assumption, however, does not necessarily hold, as enzymes of the same type can differ greatly in their kinetics^18^ and promiscuities.^19^ When this is the case, different bacteria expressing different enzyme variants will transform the contaminant at different rates, and an approach centered on microbes rather than enzymes alone may be called for to aid in the elucidation of biotransformation processes.

In this communication, we integrate metagenomics and metatranscriptomics into a genome-centric approach for identifying candidate microbes and genes that promote OMP biotransformation in complex microbial communities. The proposed approach is based on discovering relations between biotransformation rate constants and the transcriptional activities of specific microbes, as reflected by the mapping of RNA sequencing reads to metagenome-assembled genomes. Using nitrifier-rich cultures originating from activated sludge, we validate the proposed approach with ammonia, before demonstrating it on atenolol, a pharmaceutical frequently found in wastewater and removed to various degrees by biological treatment.^20^

## MATERIALS AND METHODS

### Batch experiments

Five batch experiments were conducted in total. Four batch experiments were conducted with nitrifier enrichment cultures originating from four membrane-aerated biofilm reactors inoculated with activated sludge, fed with ammonia as an electron donor, and operated each under a different aeration strategy.^21^ The fifth batch experiment was conducted with activated sludge collected at a full-scale membrane bioreactor treating hospital wastewater (Herlev, Denmark).

One volume of biomass suspended in a phosphate buffer solution was added to three volumes of a mineral medium with ammonia as an electron donor. The biomass was reactivated for 6 d in a 250-mL Erlenmeyer flask with a 65-mL working volume. The flask was loosely sealed with aluminum foil and placed on a shaker set to 120 RPM and kept at 25 °C in the dark. Following reactivation and the exhaustion of ammonia, 21 OMPs were added to a final concentration of ca. 10 µg L^-1^ each. Ammonia was spiked to a final concentration of 50 mg NH_3_-N L^-1^ (with twice the amount in moles of bicarbonate for pH control). After spiking, the flask was returned to the shaker under the same conditions.

The composition of the mineral medium was (per liter): 380 mg NH_4_Cl (100 mg NH_3_-N), 1,200 mg NaHCO_3_, 67 mg KH_2_PO_4_, 220 mg MgCl_2_•6H_2_O, 400 mg CaCl_2_•2H_2_O, 6 mg EDTA, 6 mg FeSO_4_•7H_2_O, 0.04 mg ZnSO_4_•7H_2_O, 0.08 mg CoCl_2_•6H_2_O, 0.3 mg MnCl_2_•4H_2_O, 0.2 mg CuSO_4_, 0.07 mg Na_2_MoO_4_•2H_2_O, 0.07 mg NiCl_2_•6H_2_O, 0.05 mg Na_2_SeO_4_•10H_2_O, and 0.01 Na_2_WO_4_•2H_2_O.

The added OMPs were as follows: corrosion inhibitors 1H-benzotriazole, 5-chlorobenzotriazole, and 5-methyl-1H-benzotriazole; antibiotics azithromycin, ciprofloxacin, clarithromycin, erythromycin and sulfamethoxazole; antidepressants sertraline, venlafaxine, and citalopram; beta blockers atenolol, metoprolol, and propranolol; analgesics diclofenac and ketoprofen; radiocontrast agents iohexol and iopromide; antiepileptic drug carbamazepine; immunosuppressant mycophenolic acid; and lipid-lowering agent bezafibrate, all of which often co-occur in municipal wastewater.^1,3^

### Sampling and chemical analysis

Samples for OMP and N species analyses were collected at multiple time points over 24 h after spiking. Samples for OMP analysis were mixed with acetonitrile (final concentration 20% v/v) and stored at -20 °C. Prior to analysis, the samples were thawed and filtered using 0.2 μm PTFE filters (Agilent Technologies, CA, USA). 700 µL of each filtered sample and 10 µL of an internal standard were transferred into an HPLC vial. Chromatographic separation was performed using an Eclipse C18 column installed on an Agilent 1290 Infinity HPLC system coupled with an Agilent 6470 triple quadrupole mass spectrometer (Agilent Technologies, CA, USA). The detailed HPLC-MS/MS parameters and procedures are described by Tang et al.^22^ Samples for N species analyses were filtered using 0.2 μm nylon syringe filters (Agilent Technologies, CA, USA), and ammonia, nitrite, and nitrate concentrations were determined colorimetrically with a continuous-flow auto-analyzer (Skalar, The Netherlands). Biomass samples for volatile suspended solids (VSS) measurements and DNA extraction were collected before spiking. VSS concentrations were immediately determined according to standard methods^23^ using glass microfiber filters with a pore size of 1.2 μm (Whatman, UK) and are reported in Table S1. Biomass samples for DNA extraction were centrifuged and pellets were stored at -20 °C. Biomass samples for RNA extraction were collected 5 h after spiking, snap-frozen using liquid nitrogen, and stored at -80 °C.

### Biotransformation rate constants

Ammonia and OMP removal dynamics were fitted with exponential decay curves assuming that enzyme-catalyzed biotransformation follows first-order kinetics.^5^ The first-order rate constant (*k*) was derived from the concentration versus time scatter plot using the equation C_0_e^-*k*t^, where C_0_ is the initial pollutant concentration and t is time (OriginPro 2023, OriginLab Corporation, Northampton, MA, USA) and was normalized by the VSS concentration to obtain the biotransformation rate constant (*k*_bio_).^17^ The rate constants of a pollutant across batches were compared using the Compare Datasets and Fit Parameters tool v1.7 (OriginLab Corporation, Northampton, MA, USA).

### Metagenomics and metatranscriptomics

DNA was extracted using the FastDNA SPIN Kit for soil according to the manufacturer’s protocol (MP Biomedicals, OH, USA). Shotgun DNA sequencing was performed on DNBseq (150PE; BGI, China) and yielded between 71.1-80.1 million clean reads per sample. Single-sample assembly of contigs was carried out using metaSPAdes/3.15.3^24^ with k-mer sizes 21, 33, 55, and 77. Using Vamb/3.0.3 Snakemake workflow,^25^ the contigs were binned into metagenome-assembled genomes (MAGs), which were dereplicated using dRep.^26^ GTDB-Tk/2.4.0 Classify and *de novo* workflows^27^ were respectively used to classify the dereplicated MAGs and infer their phylogenomic placement, as CoverM/0.6.1 was used to quantify them. Co-assembly of contigs was carried out using MEGAHIT/1.2.9.^28^ Contigs shorter than 1,000 bp were removed. Protein-coding genes were predicted using Prodigal/2.6.3.^29^ Total RNA extraction was accomplished using TRIzol Reagent (Invitrogen) and followed by an rRNA depletion step (TIANGEN, China). Shotgun RNA sequencing was performed on DNBseq (100PE; BGI, China) and yielded between 27.8-33.3 million clean reads per sample. The remaining rRNA sequences were removed using SortMeRNA.^30^ The transcript relative abundance was quantified using kallisto/0.46.0^31^ and is given in Transcripts Per Million (TPM). In all cases, default parameters were used, unless stated otherwise. The clean reads are available at the Sequence Read Archive (SRA) of the National Center for Biotechnology Information (NCBI) under BioProject accession number PRJNA1157596.

### Association analysis

To discover relations between biotransformation rate constants and the transcriptional activities of specific microbes, we quantified the transcript relative abundances of the genes predicted across the set of MAGs (31-43 % read alignment rate). MAGs were identified as candidate degraders (i) if their relative activity, as defined by the sum of the transcript relative abundances of the genes they carry, was positively correlated with the *k*_bio_ or (ii) if their genes were over-represented in the subset of genes whose transcript relative abundances were positively correlated with the *k*_bio_. The relative activity of a MAG and the transcript relative abundance of a gene were considered positively correlated with the *k*_bio_ if the Pearson correlation coefficient was positive and different from zero at a significance level of 0.01. The over-representation analysis was performed using the enricher function of the clusterProfiler R package with the Benjamini-Hochberg procedure for *p* value adjustment.^32^ No correction for multiple hypothesis testing was applied to the significance levels of the correlations between the relative activities of the MAGs and the biotransformation rate constants because of the small number of data points with relation to the number of MAGs. A step-by-step depiction of the two methods used to identify candidate degraders is provided by Figure S1.

### Expression of genes at the community level

To find evidence supporting our genome-centric association analysis, we examined the expression of genes that are associated with the biotransformation of specific pollutants at the community level. To this end, we quantified the transcript relative abundances of the genes predicted across the set of the co-assembled contigs (50-55 % read alignment rate). The expression of a gene of interest at the community level was then calculated by summing the transcript relative abundances of all the identified instances of the gene in this set.

### Gene annotation

Genes were searched using hmmsearch with hidden Markov model (HMM) profiles as downloaded from the NCBI Protein Family Models database and default parameters (HMMER 3.3.2; http://hmmer.org/). The minimum hmmsearch full-sequence score for the proteins to be considered hits were the sequence cutoffs as they appear in the database.

## RESULTS AND DISCUSSION

### Pollutant removal dynamics

Ammonia removal followed first-order kinetics (*R*^2^ = 0.98 ± 0.01; Figure S2 and Table S1) with statistically different rate constants across batches (*F* test, *p* value < 0.0001). All the ammonia was oxidized to nitrite or nitrate and no denitrification was detected (Figure S3). Among the added OMPs, the removal of atenolol followed first-order kinetics (*R*^2^ = 0.89 ± 0.14; Figure S4 and Table S1) with statistically different rate constants across batches (*F* test, *p* value < 0.0001). Its biotransformation rate constants ranged between 3-12 L gVSS^-1^ d^-1^ and resembled the biotransformation rate constants of atenolol previously reported for activated sludge or moving bed biofilm reactors.^33^ Mycophenolic acid was rapidly eliminated by the activated sludge in a pattern matching first-order kinetics (*R*^2^ = 0.99; Figure S5), like in an earlier report. ^22^ However, its removal by the nitrifier enrichments poorly matched this type of kinetics (*R*^2^ = 0.44 ± 0.38). Sertraline was also eliminated by the activated sludge, but its concentration leveled out in batches with the nitrifier enrichments after an initial drop (Figure S5), a pattern previously attributed to sorption of the antidepressant.^22^ Bezafibrate was removed at an efficiency of between 25 to 60 % (Figure S5). However, while its removal followed first-order kinetics in all the batches (*R*^2^ = 0.88 ± 0.07), the rate constants were not statistically different across the different biomass types (*F* test, *p* value = 0.2). The other OMPs were persistent (removal efficiency < 30 %) even though some of them, such as clarithromycin, diclofenac, iohexol, or ketoprofen, were removed in other experimental aerobic reactors, which were fed with wastewater.^5,22^ Therefore, we focused the association analysis on ammonia and atenolol, using the former pollutant for validation.

### Microbial community composition

In total, 159 metagenome-assembled genomes (MAGs) were retrieved, to which 67 % of the DNA reads were recruited on average (SD 20; Figure S6). Details on the MAGs, including their classifications and relative abundances, can be found in Table S2. Based on MAG classification and DNA read coverage, *Pseudomonadota* and *Bacteroidota* dominated all microbial communities. *Gemmatimonadota* were only present in the nitrifier enrichments, while *Acidobacteriota* and *Actinomycetota* were more abundant in the activated sludge (Figure S7). *Nitrosomonadaceae*, which comprises ammonia-oxidizing bacteria (AOB), was the most abundant family in the nitrifier enrichments with relative abundances of between 25-43 % and the fourth most abundant family in the activated sludge with a relative abundance of 9 % (Figure S8). It was composed of *Nitrosomonas europaea* (MAG S1C1150), found in high numbers in all the five batches, *Nitrosomonas mobilis* (MAG S2C1424), found in low numbers in the nitrifier enrichments, and *Nitrosospira* sp. (MAG S2C2712), found in high numbers especially in two of the nitrifier enrichments (Figure S9), in agreement with the cultures of origin. ^21^

### Association analysis: ammonia as validation

The *k*_bio_ of ammonia was positively correlated with the relative activity of MAG S1C1150 (Pearson *r* = 0.98; Figure 1A and Table S3), which was classified as the ammonia oxidizer *N. europaea*. It was not positively correlated with the relative activity of any other MAG (Table S3). S1C1150 was also the MAG whose genes were the most over-represented among the genes whose transcript relative abundances were positively correlated with the *k*_bio_ of ammonia. Specifically, the transcript relative abundance of 246 genes correlated with the *k*_bio_ of ammonia, of which 194 belonged to MAG S1C1150, while no more than 8 belonged to any other MAG (Table S4). The distribution of the coefficient of the correlation between the *k*_bio_ of ammonia and the transcript relative abundances of all the genes in MAG S1C1150 had a strong negative skew with Pearson *r* > 0.8 in 90 % of the genes (Figure 1B). This skew shows that the removal of ammonia was not correlated only with the expression of a particular gene or a subset of genes that belong to a particular pathway but rather with the transcriptional activity of *N. Europaea* as a whole.

**Figure 1.**
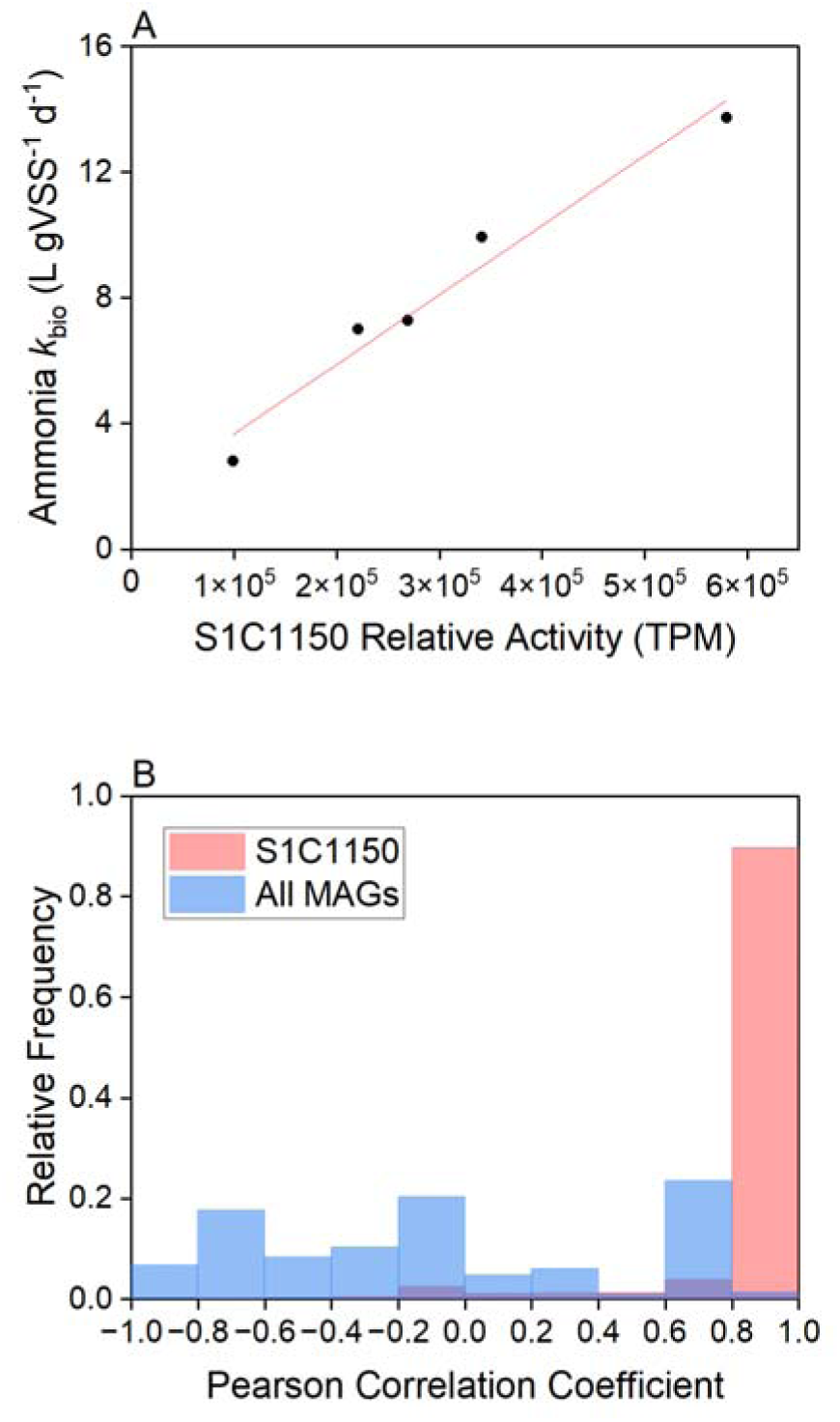
(A) The biomass-normalized first-order rate constant (*k*_bio_) of ammonia as a function of the relative activity of MAG S1C1150 (*N. europaea*). The relative activity was calculated by summing the transcript relative abundances of all the genes in the MAG. The red line is a linear fit (Pearson *r* = 0.98, *p* value = 0.003). (B) Overlaying histograms representing the distributions of the Pearson correlation coefficient between the gene transcript relative abundances and the *k*_bio_ of ammonia for the genes in MAG S1C1150 (light red) and in all the MAGs (light blue).

### Association analysis: atenolol

The *k*_bio_ of atenolol positively correlated with the relative activity of a single MAG, namely S4C147, which was classified as a *Bacteroidota* bacterium of the genus *Terrimonas* (Pearson *r* = 1.00; Figure 2A). The most over-represented genes among the genes whose transcript relative abundance positively correlated with the *k*_bio_ of atenolol belonged to MAG S4C5535, which was classified as a *Pseudomonadota* bacterium of the genus *Zeimonas* (hypergeometric test, over-representation *p* value < 10^-300^; Table S5). As before, the distribution of the coefficient of the correlation between the *k*_bio_ of atenolol and the transcript relative abundances of the genes carried by S4C5535 was negatively skewed with Pearson *r* > 0.8 in 62 % of the genes (Figure 2B). Besides S4C5535, several other MAGs could be associated with atenolol biotransformation according to the over-representation analysis. These MAGs included S3C5076, S4C147, and S1C1358, to which the next most over-represented genes belonged (hypergeometric test, over-representation *p* values < 10^-90^; Table S5 and Figure S10). Like S4C147, S3C5076 was classified to the genus level as *Terrimonas*, while S1C1358 was classified to the same level as *Flavobacterium*. The over-representation analysis results for atenolol are depicted in Figure 3. As Figure 3 and Table S3 show, S4C5535, S3C5076, S4C147, and S1C1358 were not among the most active MAGs, having a low to medium relative activity.

**Figure 2.**
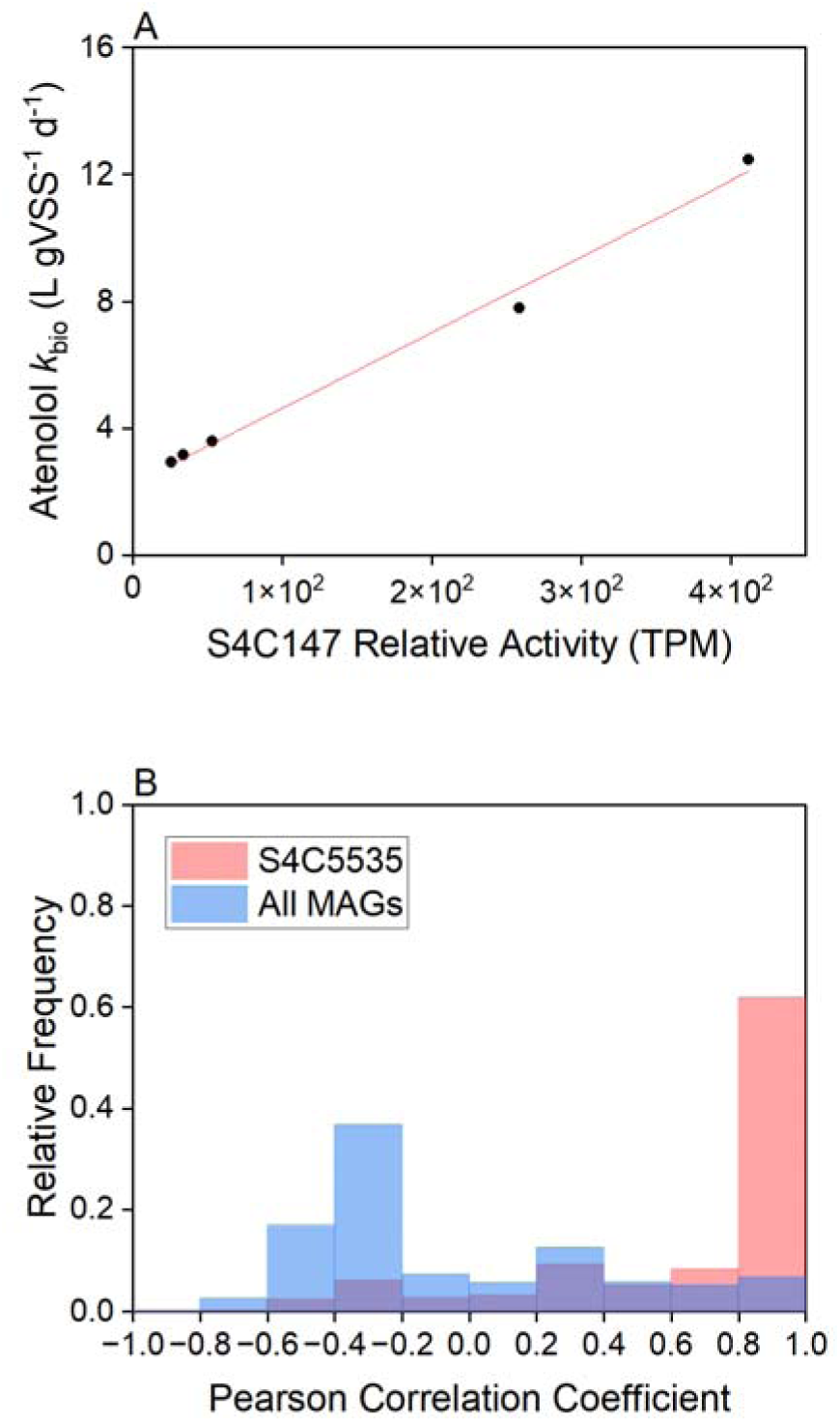
(A) The biomass-normalized first-order rate constant (*k*_bio_) of atenolol as a function of the relative activity of MAG S4C147 (*Terrimonas* sp.). The relative activity was calculated by summing the transcript relative abundances of all the genes in the MAG. The red line is a linear fit (Pearson *r* = 1.00, *p* value = 0.0003). (B) Overlaying histograms representing the distributions of the Pearson correlation coefficient between the gene transcript relative abundances and the *k*_bio_ of atenolol for the genes in MAG S4C5535 (*Zeimonas* sp.; light red) and in all the MAGs (light blue).

**Figure 3.**
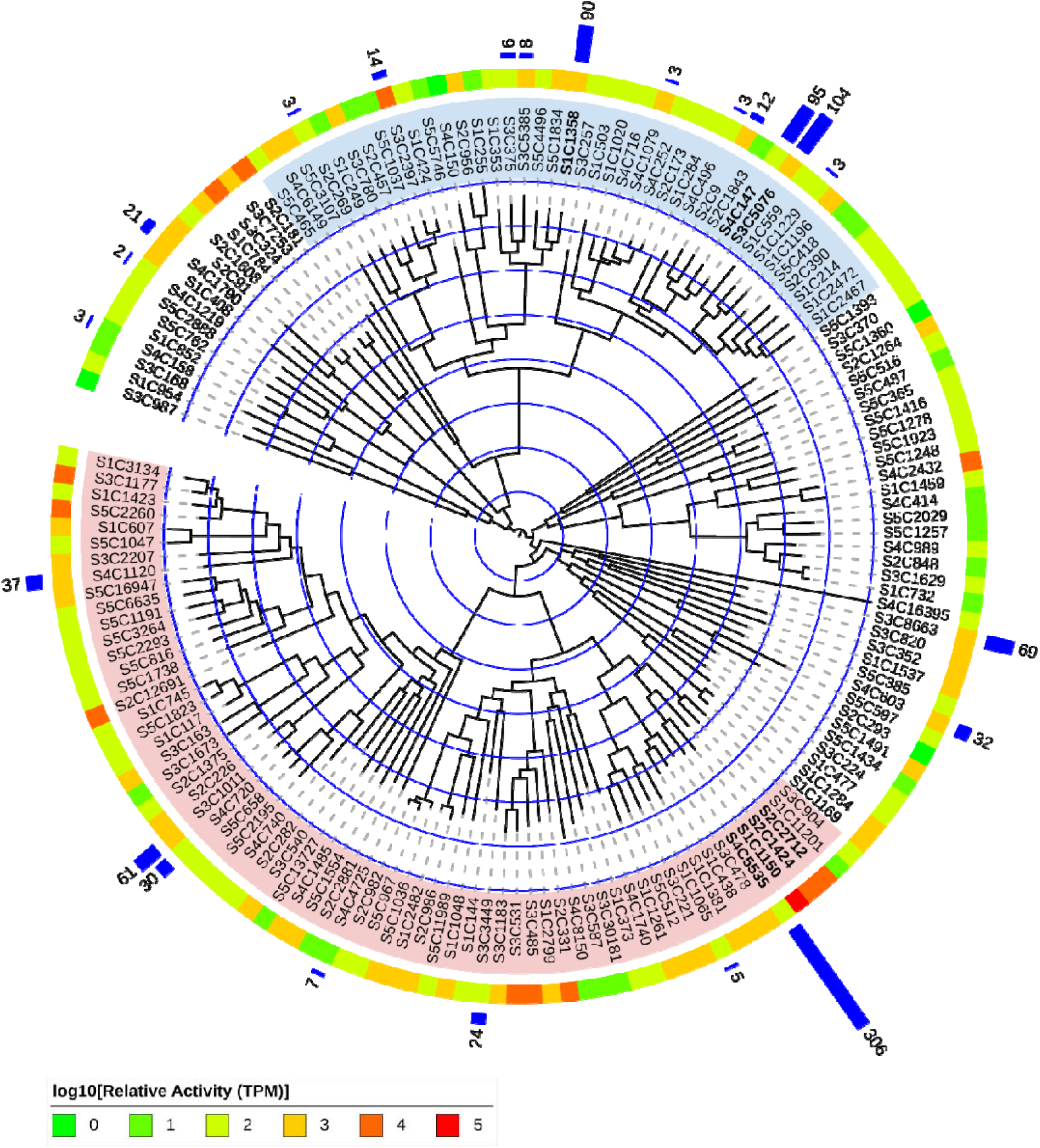
Phylogenomic tree of the MAGs with their average relative activity (log10; color gradient) and the p value for the over-representation of their genes in the subset of genes whose transcript relative abundances were positively correlated with the *k*_bio_ of atenolol (-log10; blue bars). The average relative activity is the average of the relative activities in all batches. The relative activity in each batch was calculated by summing the transcript relative abundances of all the genes in the MAG, which are given in units of transcripts per million (TPM). The *p* value was calculated based on the hypergeometric distribution and was corrected for multiple hypothesis testing using the Benjamini-Hochberg procedure. The *Pseudomonadota* and *Bacteroidota* are indicated by light red and light blue, respectively. The MAGs discussed in the text are highlighted in bold.

*Zeimonas*, *Terrimonas*, and *Flavobacterium* are heterotrophic bacteria.^34^ Indeed, atenolol biotransformation by nitrifying microbiomes was attributed to heterotrophic members of the communities.^13,15,21^ *Terrimonas* and *Flavobacterium* are commonly found in activated sludge and belong to the phylum *Bacteroidota*.^34^ Evidence that *Bacteroidota* consume products produced by AOB^10^ and that atenolol biotransformation is negatively affected by nitrification inhibition^15^ jointly support the possibility of the two genera being atenolol degraders. In particular, the relative abundance of *Terrimonas* and the biotransformation rate constant of atenolol increased in a hybrid biofilm reactor for methane valorization following an increase of the methane loading rate.^35^

The advantage of the over-representation analysis was that MAGs met the threshold for statistical significance after correcting for multiple hypothesis testing. The correlations between the relative activities of the MAGs and the biotransformation rate constants of ammonia and atenolol could not meet such a threshold because of the small number of data points with relation to the number of hypotheses tested, as previously recognized in similar association studies with omics-scale data.^16,17^ Nonetheless, the successful application of the relative activity-based method and the overrepresentation-based method to ammonia suggests that the identifications of degraders made by both methods are valid.

### Correlations between pollutant biotransformations and targeted enzymes

We examined the correlations between the biotransformation rate constants of ammonia and atenolol and the transcript relative abundances of genes that are associated or putatively associated with their biotransformation.

AOB oxidize ammonia with the ammonia monooxygenase (AMO) enzyme. AMO catalyzes the first step of nitrification by converting ammonia to hydroxylamine and comprises three subunits: AmoA, AmoB, and AmoC.^36^ We found no correlation between the *k*_bio_ of ammonia and the expression of AMO at the community level, calculated as the sum of the transcript relative abundances of the genes identified as *amoA* (NCBI HMM accession NF041557.1), *amoB* (NCBI HMM accession NF041640.1) or *amoC* (NCBI HMM accession NF041641.1) in the set of co-assembled contigs (Pearson *r* = 0.48, *p* value = 0.41). On the other hand, we found a positive correlation between the *k*_bio_ of ammonia and the expression of AMO calculated as the sum of the transcript relative abundances of the *amoA*, *amoB*, and *amoC* genes of *N. europaea* (Pearson *r* = 0.94, *p* value = 0.02). These findings support the premise of our genome-centric approach, according to which the transcript relative abundance of a single enzyme within a set of enzymes with the same specific function may predict the *k*_bio_ of the substrate. We speculate that the different AMOs variants have different reaction kinetic parameters. Such variability can explain our results and, in the wider sense, the documented different niche preferences and substrate affinities of *N. europaea*, *N. mobilis*, and *Nitrosospira* spp.^21,37,38^ The *amo* genes were largely absent from the MAGs classified as AOB, a common issue with these genes, which are multi-copy.^39^ They were assigned to a genus by phylogenetic placement (Figure S11).

Atenolol biotransformation in activated sludge has been shown to be initiated by primary amide hydrolysis, which is catalyzed by amidohydrolases.^40,41^ We tested the *k*_bio_ of atenolol against the expression of a comprehensive set of 20 primary amide amidohydrolases, compiled based on KEGG REACTION database entries RC00010 and RC02798.^42^ No significant correlation was found between the biotransformation rate constant of atenolol and the expression of the amidohydrolases at the community level, whether they were considered separately or collectively (Table S6). This observation implies that atenolol may be transformed by a specific variant of a specific amidohydrolase, in line with our premise.

Indeed, amidohydrolases that share the same primary function can differ in their Michaelis constants and maximum velocities^18^ and in their promiscuous activities.^19^ Therefore, the contribution of each of such amidohydrolases to atenolol biotransformation may not necessarily be proportional to its transcript relative abundance and so the rate of atenolol removal could be determined by a subset of cell types. We found 11 primary amide amidohydrolases in MAGs S4C147, S3C5076, S4C5535, or S1C1358, ten of which were glutamine-hydrolyzing (Table S6). The transcript relative abundance of asparagine synthase (AsnB), which catalyzes the transfer of the amide nitrogen of glutamine to aspartate, was positively correlated with the *k*_bio_ of atenolol in the cases of MAGs S4C147, S3C5076, and S4C5535 (Table S7).

In contrast to our results, Achermann *et al.* found the removal rate of ammonia to be correlated with the total relative abundance of *amo* transcripts at the community level.^17^ Also in contrast to our results, Johnson *et al.* discovered through association analysis a positive, monotonic, albeit not linear correlation among ten wastewater treatment plant communities between the biotransformation rate constant of atenolol and the total relative abundance of the transcripts encoding the amidohydrolase urease,^16^ which was included in our analysis (Table S6). Therefore, the enzyme-centric approach, which these two studies represent, and the genome-centric approach, which we propose here, can be considered complementary to one another. Compared with the enzyme-centric approach, the genome-centric approach compensates for reaction kinetics variability and can consequently prove to be useful in identifying OMP biotransformation drivers when multiple microorganisms capable of performing the same metabolic function coexist.

### Implications

We introduce an association analysis-based method for identifying candidate microbes and enzymes that set the biotransformation rates of OMPs in complex microbial communities. Being robust to variability in enzyme kinetics across microbial species, the method complements previously introduced association analysis-based methods for identifying candidate enzymes that transform OMPs.^16,17^ Correlations between the removal of OMPs and the activity of specific microbes and genes can allow for the prediction of OMP fate in natural and engineered environments. Furthermore, they can inform studies designed to establish causality between microbes or enzymes and the removal of OMPs using, for example, pure cultures or cells genetically modified to express specific enzymes, as well as metabolic modeling efforts seeking to enhance OMP removal. We specifically identified candidate bacteria and genes that are involved in the biotransformation of atenolol and whose activity or expression can potentially be used to assess the capacity of microbiomes to remove this widespread OMP.

## ASSOCIATED CONTENT

### Supporting Information

Supporting figures: pollutant removal dynamics, DNA alignment rate, microbial community composition, change point detection, and AMO phylogenetic placements (.pdf)

Supporting tables: additional information on rate constants, MAGs, and the association analyses results (.xlsx)

### Notes

The authors declare no competing financial interest.

## ACKNOWLEDGEMENTS

The research leading to these results was supported by the Novo Nordisk Foundation with grant NNF21OC0071581 (VIRTUE) to B.F.S. B.V-P acknowledges the financial support provided by the Danish Research Council for Independent Research under the Sapere Aude DFF Starting Grant 10.46540/2067–00029B.

