## Supplementary figures for "An integrated meta-omics approach for identifying candidate organic micropollutant degraders in complex microbial communities"

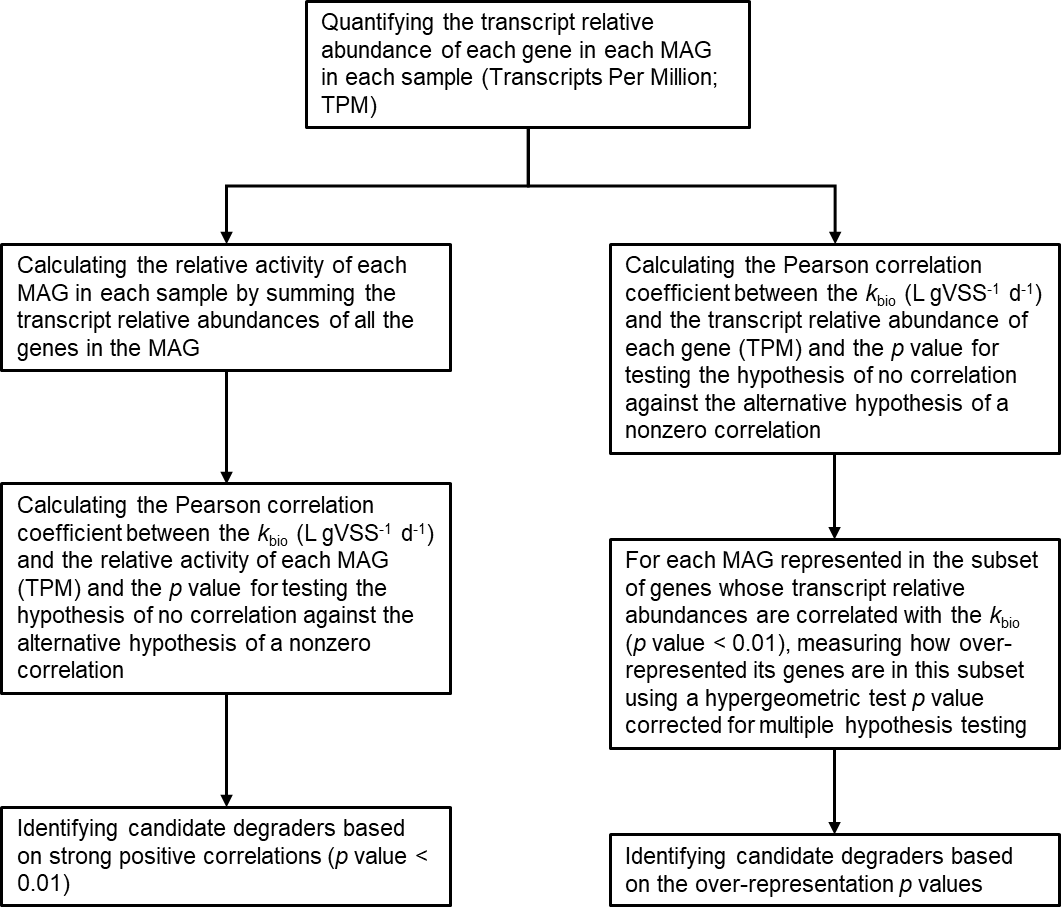

### **Figure S1.** A step-by-step depiction of the two methods used to identify candidate degraders.

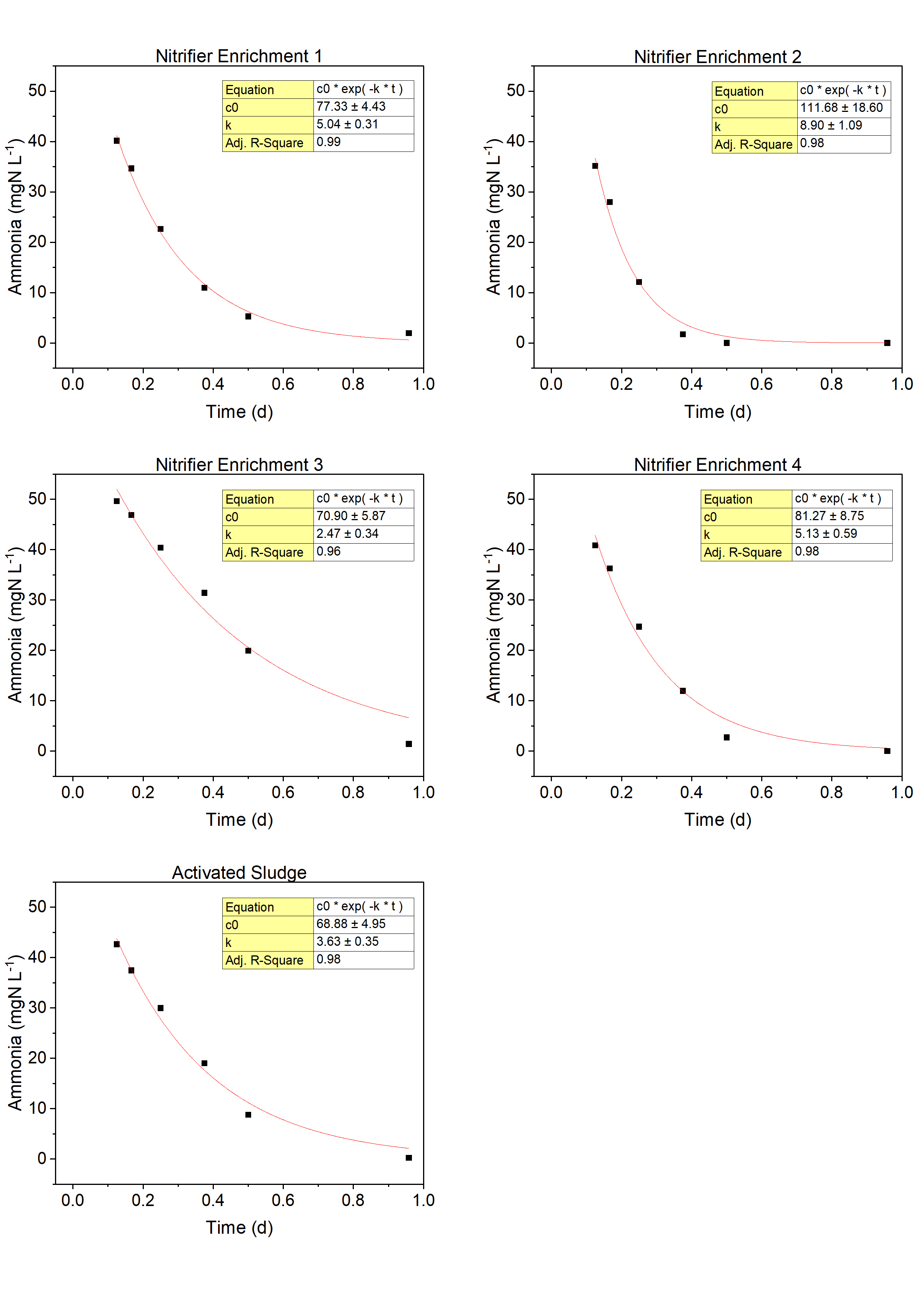

### **Figure S2.** Ammonia concentration as a function of time and exponential decay fits.

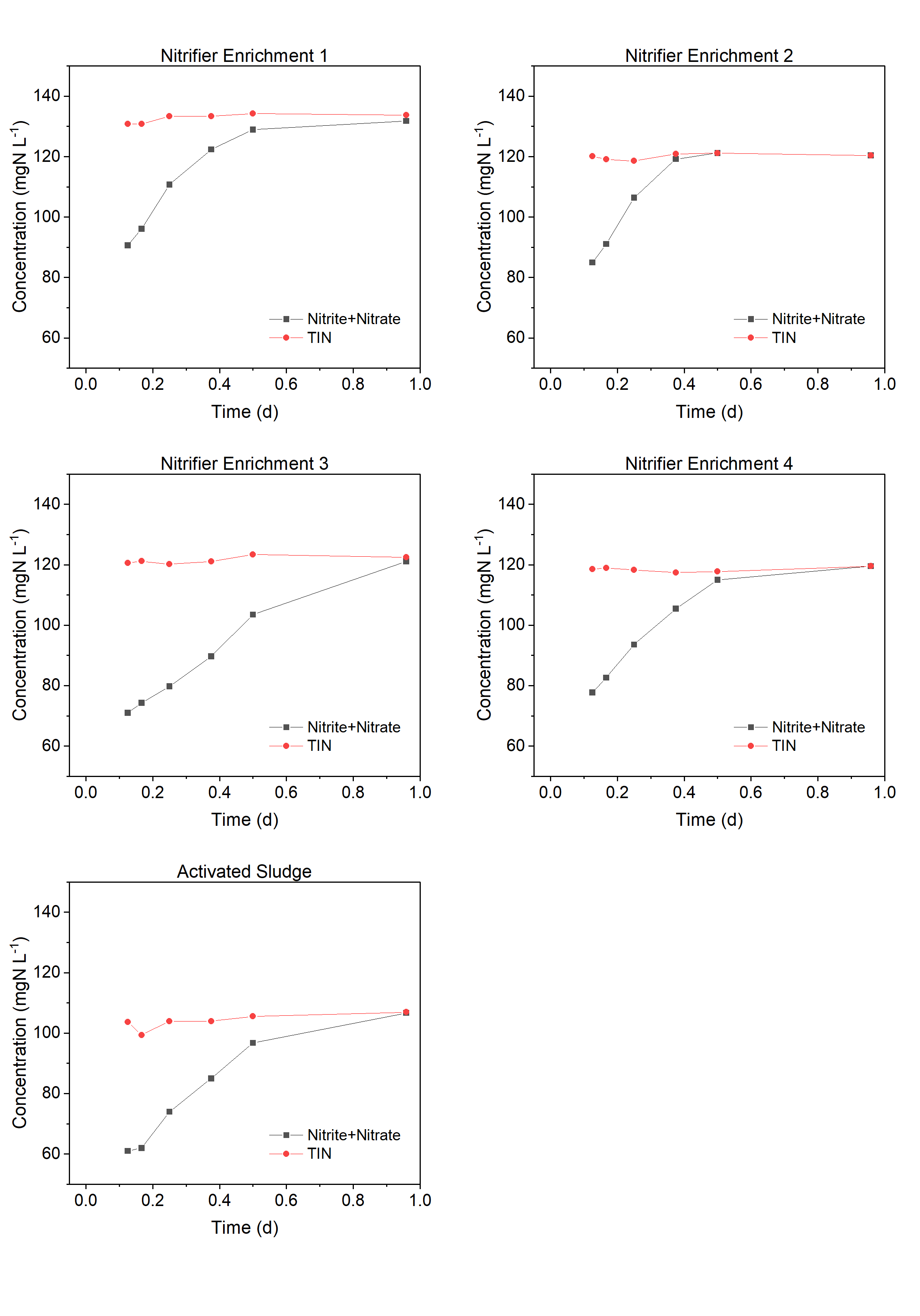

### **Figure S3.** Concentrations of Nitrite+nitrate and Total Inorganic Nitrogen (ammonia+nitrite+nitrate; TIN) as a function of time.

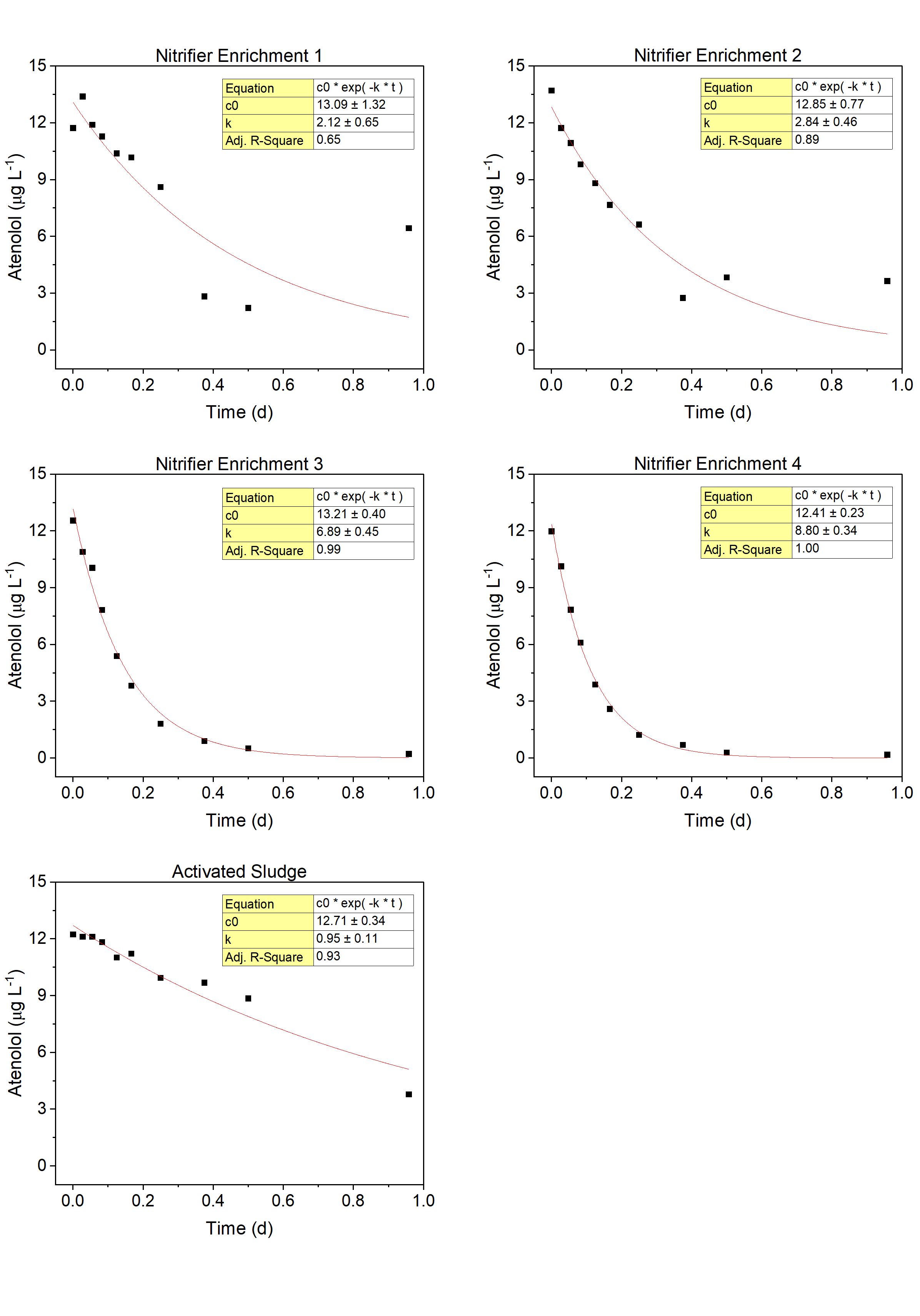

### **Figure S4.** Atenolol concentration as a function of time and exponential decay fits.

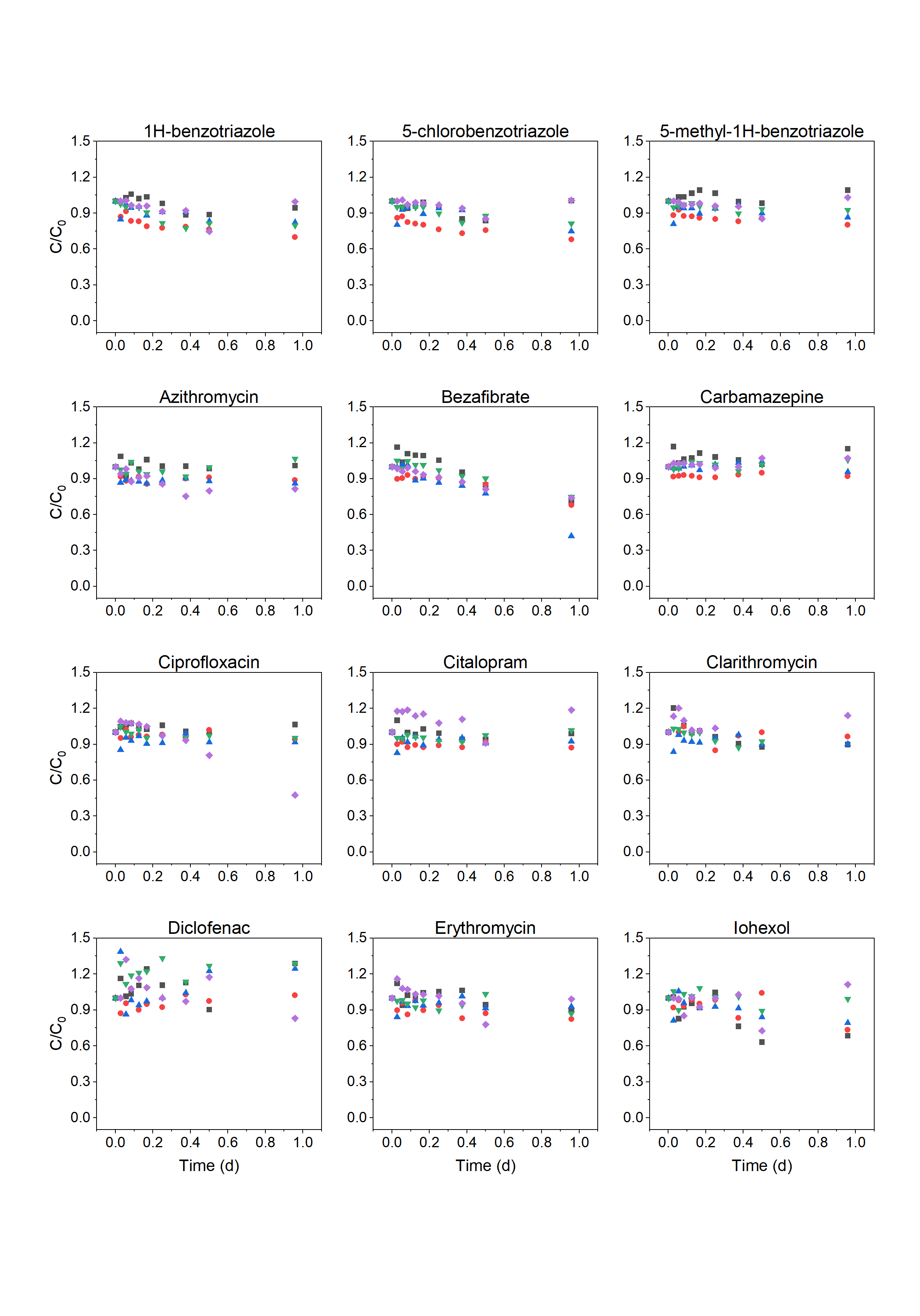

### **Figure S5.** OMP removal dynamics. Black triangles – Nitrifier Enrichment 1; Red circles – Nitrifier Enrichment 2; Blue up-pointing triangles – Nitrifier Enrichment 3; Green down-pointing triangles – Nitrifier Enrichment 4; Purple diamonds – Activated Sludge. C/C_0_ denotes the concentration normalized to the initial concentration. Values greater than 1 are due to measurement uncertainty.

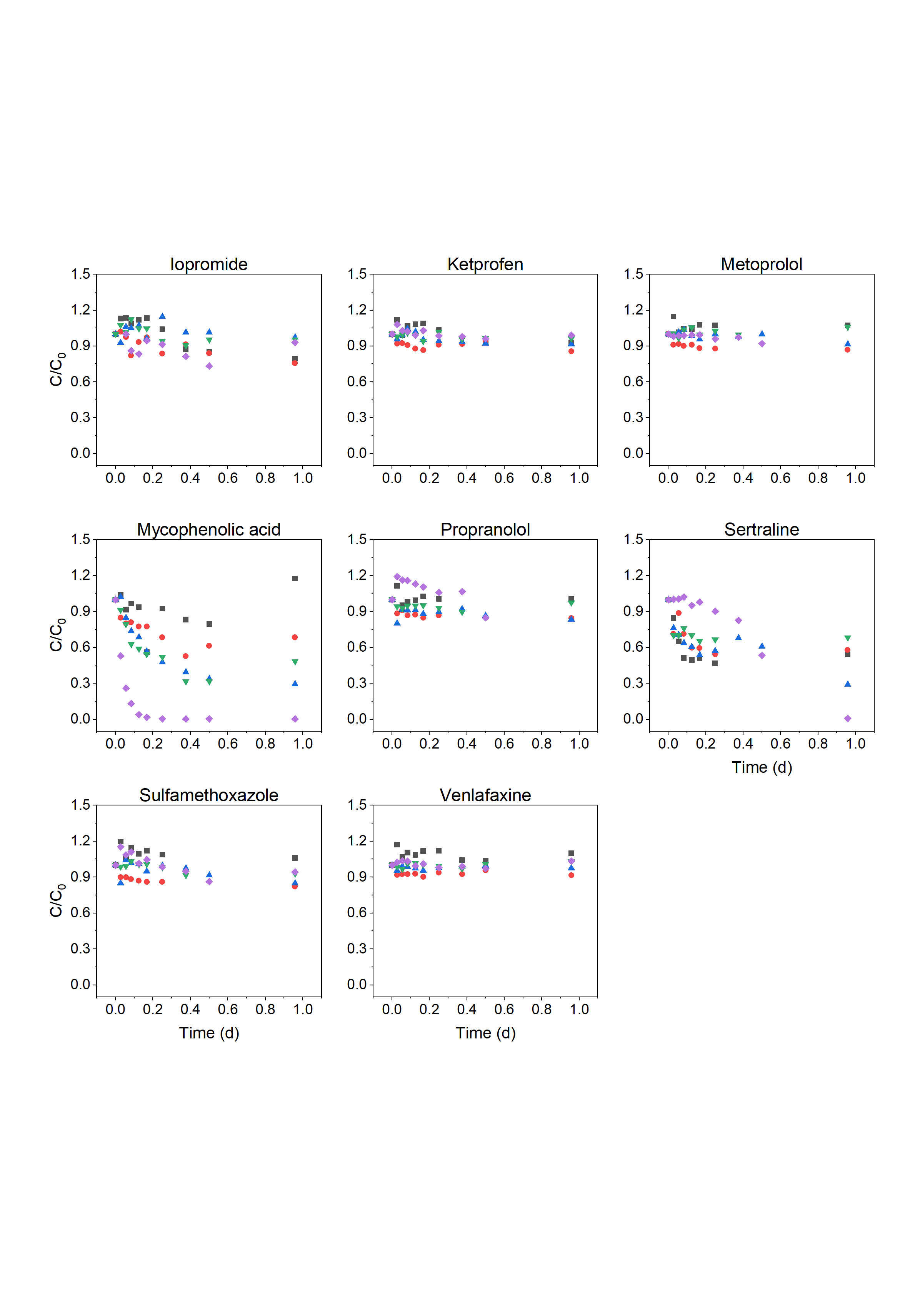

### **Figure S5 (continued).** Black triangles – Nitrifier Enrichment 1; Red circles – Nitrifier Enrichment 2; Blue up-pointing triangles – Nitrifier Enrichment 3; Green down-pointing triangles – Nitrifier Enrichment 4; Purple diamonds – Activated Sludge. C/C_0_ denotes the concentration normalized to the initial concentration. Values greater than 1 are due to measurement uncertainty.

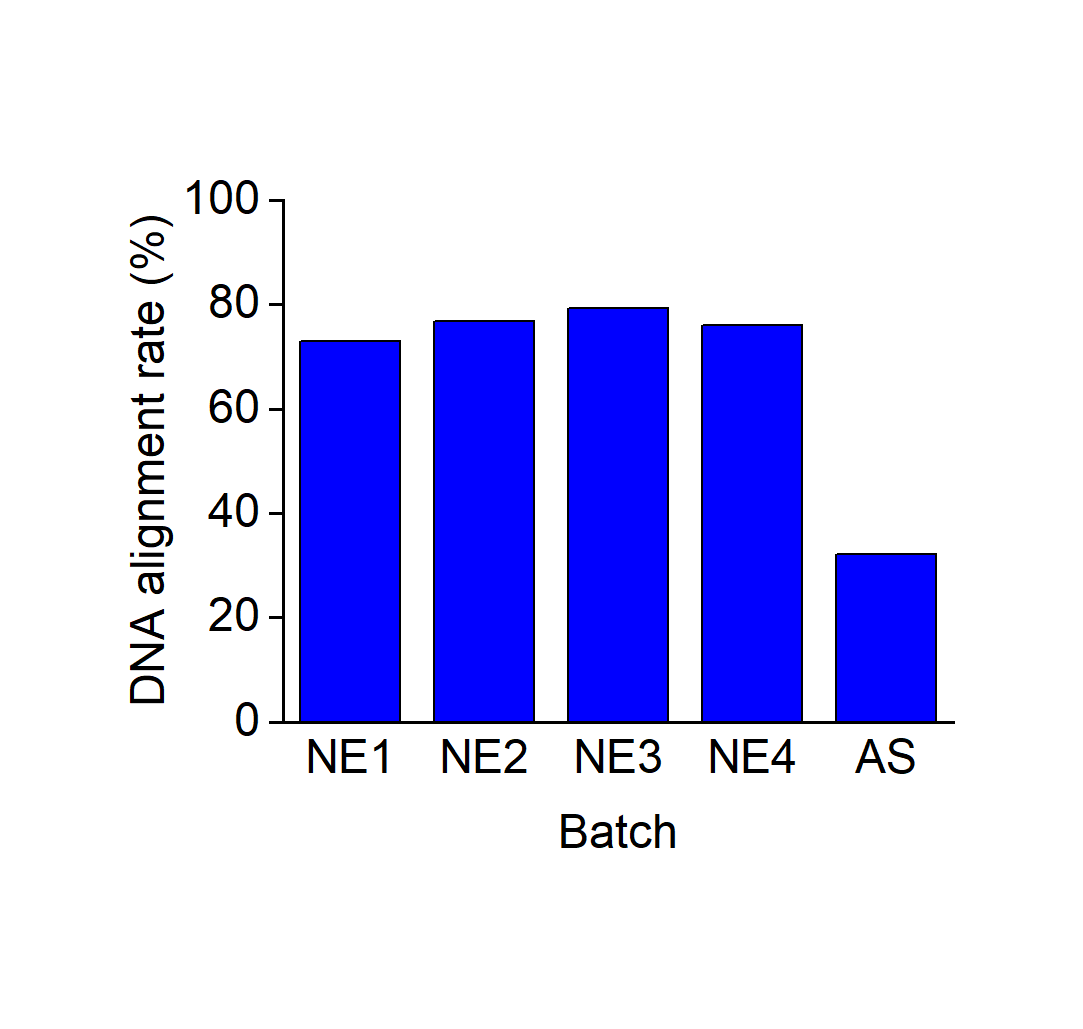

### **Figure S6.** DNA alignment rate. The percentage of DNA reads that was mapped to the retrieved MAGs. NE: Nitrifier Enrichment; AS: Activated Sludge.

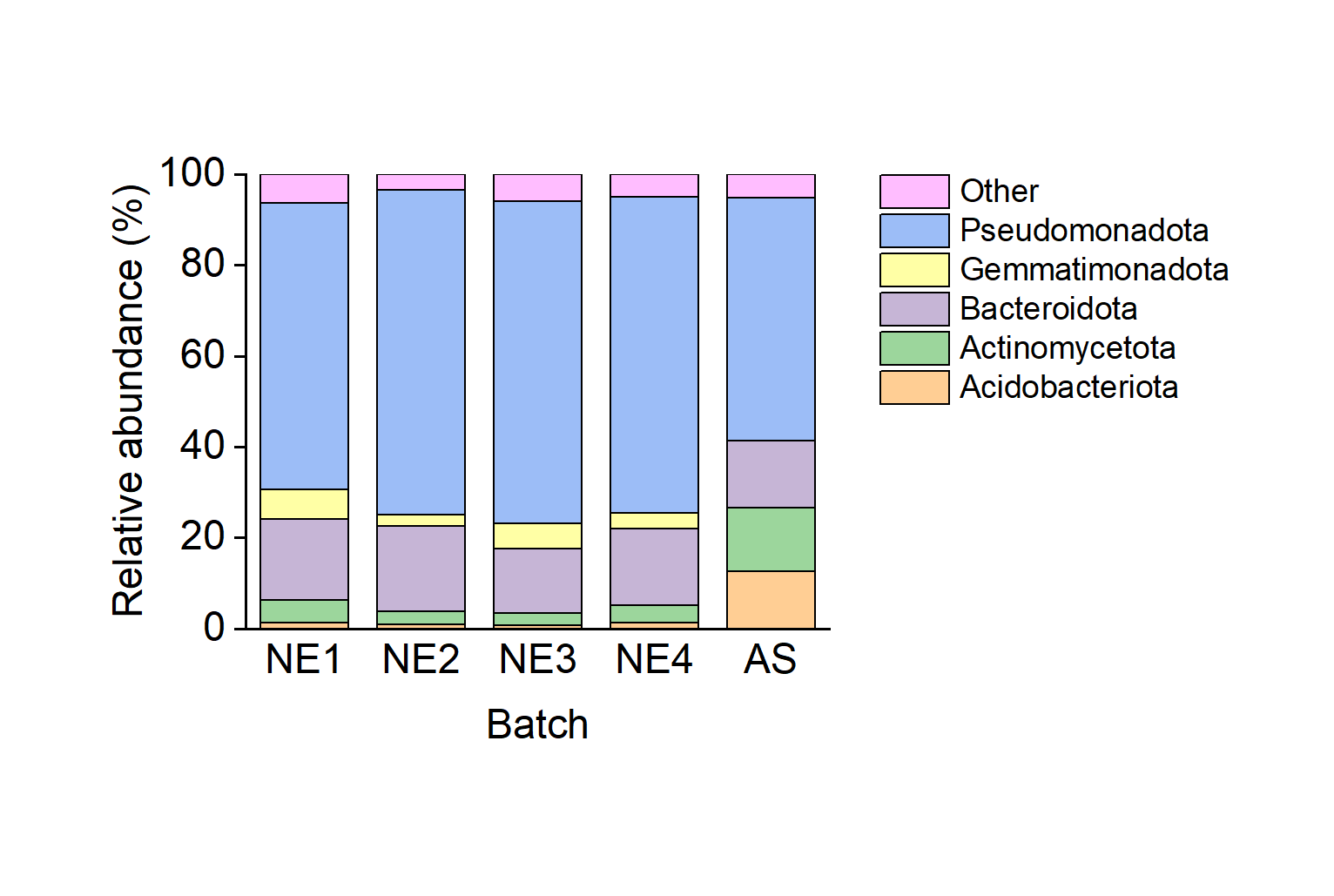

### **Figure S7.** Microbial community composition at the phylum level based on the classification and the DNA read coverage of the MAGs. Shown are phyla with a relative abundance greater than 2 % in at least one of the batches. NE: Nitrifier Enrichment; AS: Activated Sludge.

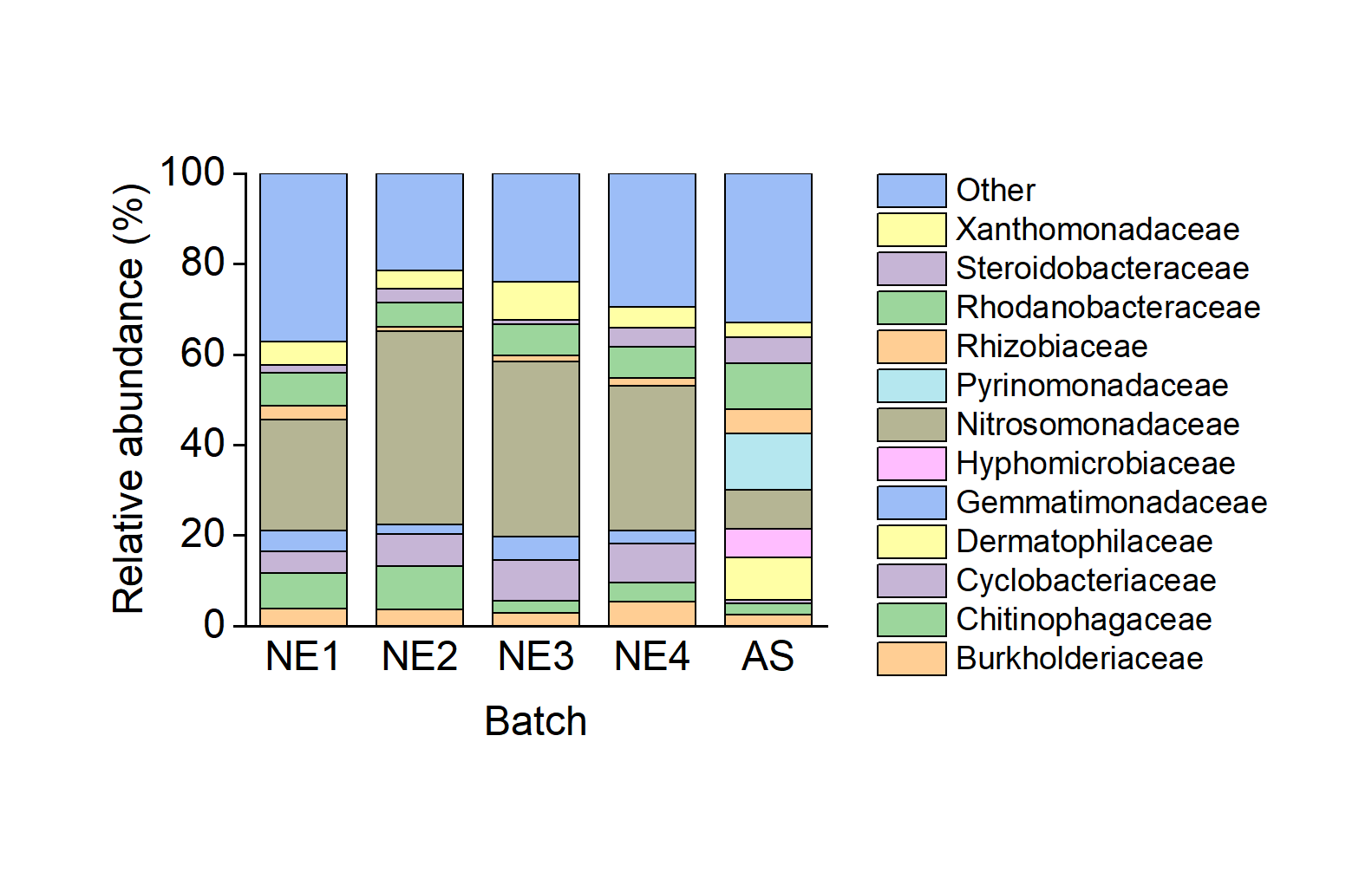

### **Figure S8.** Microbial community composition at the family level based on the classification and the DNA read coverage of the MAGs. Shown are families with a relative abundance greater than 5 % in at least one of the batches. NE: Nitrifier Enrichment; AS: Activated Sludge.

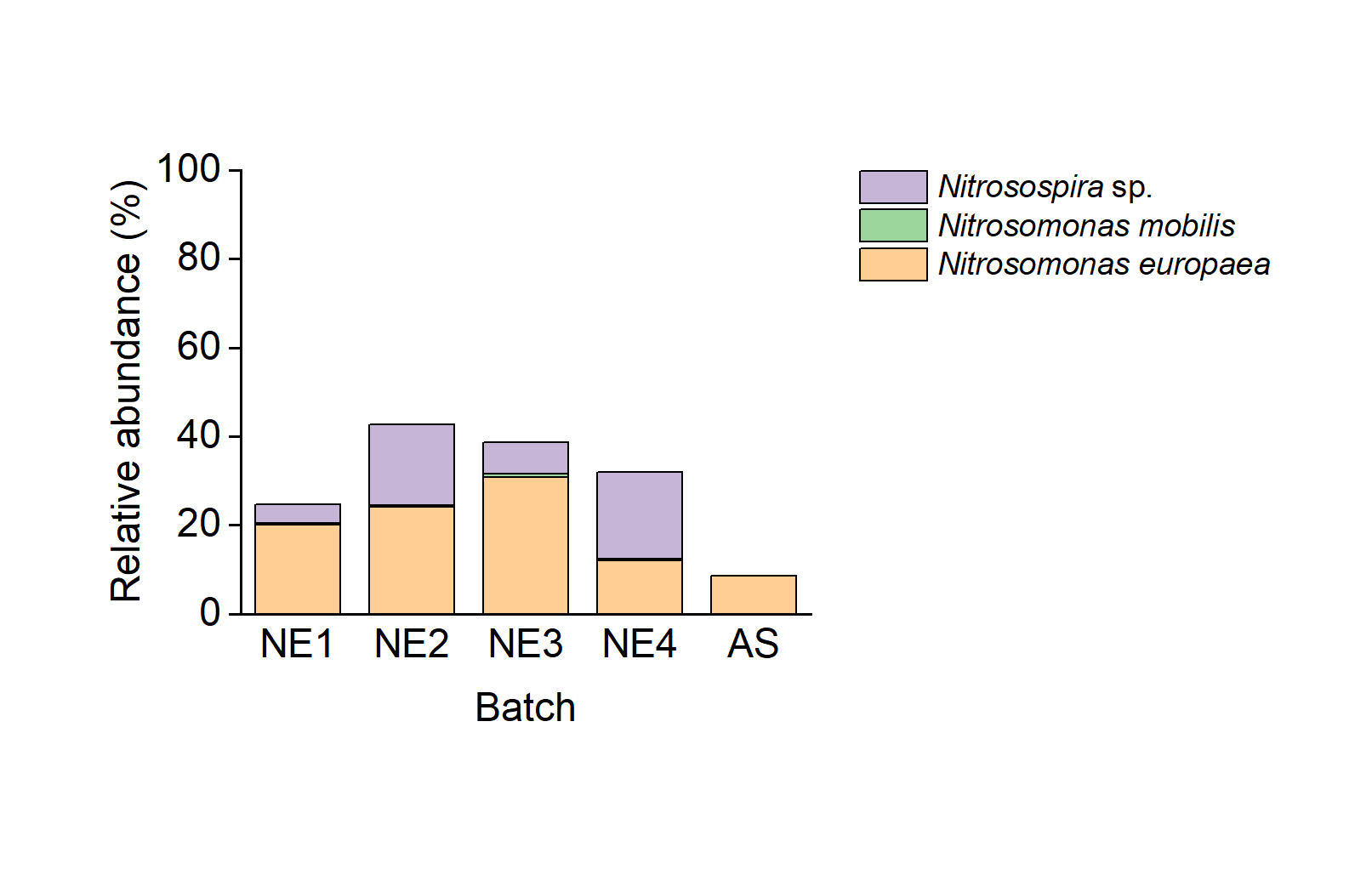

### **Figure S9.** AOB relative abundance based on the classification and the DNA read coverage of the MAGs. NE: Nitrifier Enrichment; AS: Activated Sludge.

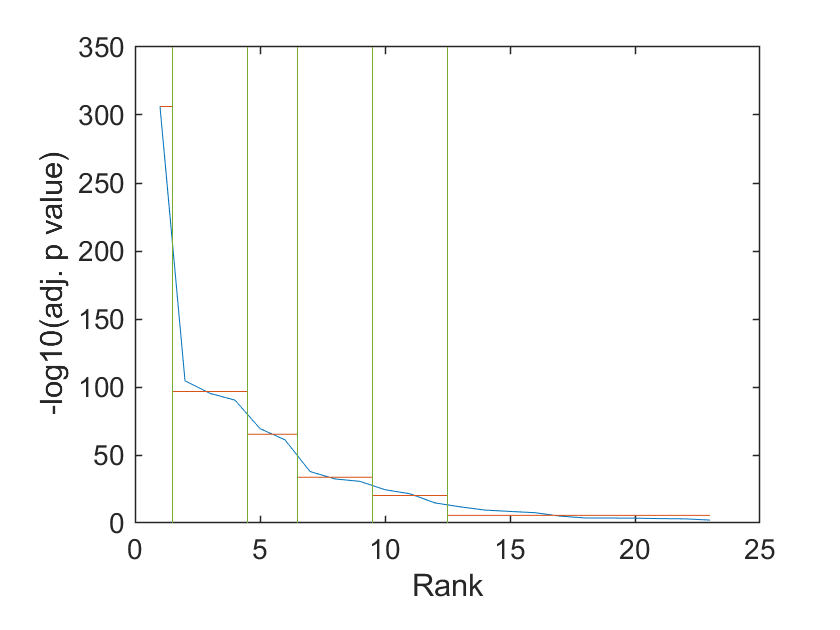

### **Figure S10.** Change points in the adjusted *p* value of the over-representation analysis for atenolol. The genes of 23 MAGs were over-represented (adjusted *p* value < 0.05) in the subset of genes whose transcript relative abundances were correlated with the biotransformation rate constant of atenolol (Table S5). The 23 adjusted *p* values are plotted as a function of their rank. The vertical lines represent the points at which the adjusted *p* value changes most significantly. The change points were obtained by minimizing the sum of the residual errors of all the regions from their local means.

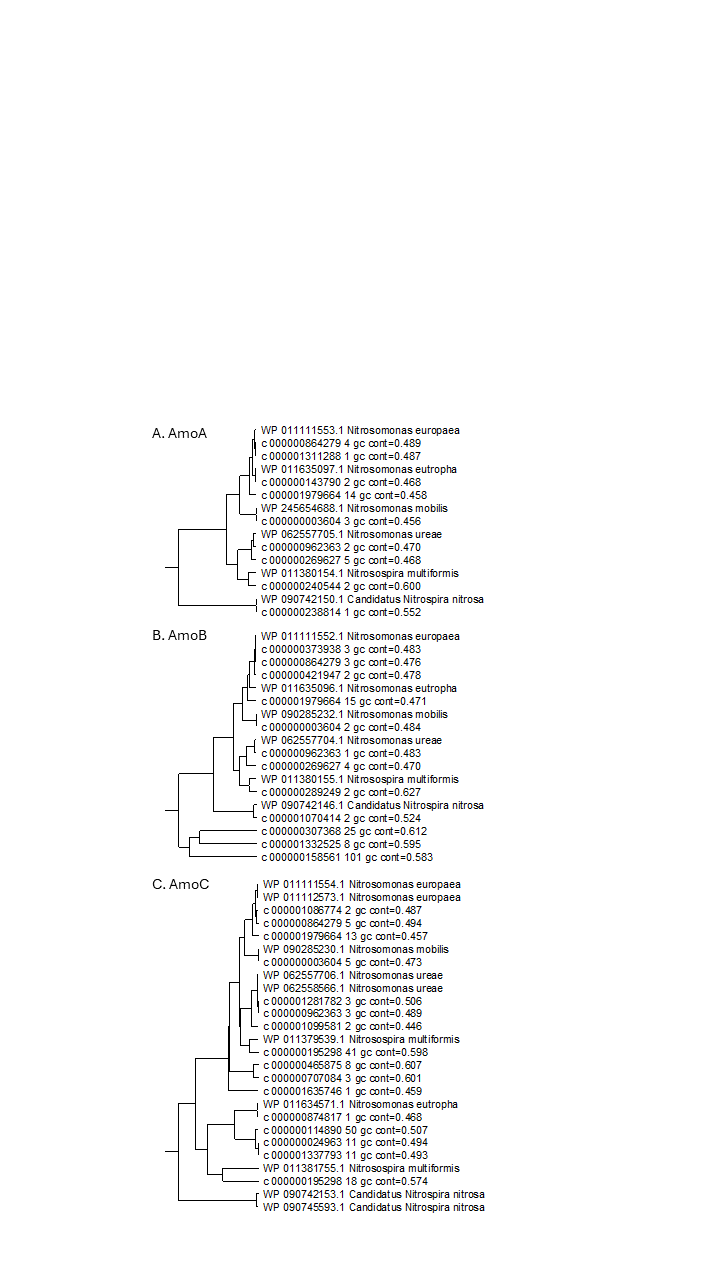

### **Figure S11.** Phylogenetic placements for AmoA (A), AmoB (B), and AmoC (C). Sequences with the prefix ‘c’ represent predicted protein sequences annotated using HMM profiles. Sequences with the prefix ‘WP’ represent (non-redundant) RefSeq protein sequences.
